# Automatic navigation system for the mouse brain

**DOI:** 10.1101/442558

**Authors:** Susan J. Tappan, Brian S. Eastwood, Nathan O’Connor, Quanxin Wang, Lydia Ng, David Feng, Bryan M. Hooks, Charles R. Gerfen, Patrick R. Hof, Christoph Schmitz, Jack R. Glaser

## Abstract

Identification and delineation of brain regions in histologic mouse brain sections is especially pivotal for many neurogenomics, transcriptomics, proteomics and connectomics studies, yet this process is prone to observer error and bias. Here we present a novel brain navigation system, named NeuroInfo, whose general principle is similar to that of a global positioning system (GPS) in a car. NeuroInfo automatically navigates an investigator through the complex microscopic anatomy of histologic sections of mouse brains (thereafter: “experimental mouse brain sections”). This is achieved by automatically registering a digital image of an experimental mouse brain section with a three-dimensional (3D) digital mouse brain atlas that is essentially based on the third version of the Allen Mouse Brain Common Coordinate Framework (CCF v3), retrieving graphical region delineations and annotations from the 3D digital mouse brain atlas, and superimposing this information onto the digital image of the experimental mouse brain section on a computer screen. By doing so, NeuroInfo helps in solving the long-standing problem faced by researchers investigating experimental mouse brain sections under a light microscope—that of correctly identifying the distinct brain regions contained within the experimental mouse brain sections. Specifically, NeuroInfo provides an intuitive, readily-available computer microscopy tool to enhance researchers’ ability to correctly identify specific brain regions in experimental mouse brain sections. Extensive validation studies of NeuroInfo demonstrated that this novel technology performs remarkably well in accurately delineating regions that are large and/or located in the dorsal parts of mouse brains, independent on whether the sections were imaged with fluorescence or brightfield microscopy. This novel navigation system provides a highly efficient way for registering a digital image of an experimental mouse brain section with the 3D digital mouse brain atlas in a minute and accurate delineation of the image in real-time.

## INTRODUCTION

Microscopic analysis of histologic brain sections provides important information about the organization and structure of the nervous system, especially important for neurogenomics, transcriptomics, proteomics and connectomics studies. In this context, the BRAIN Initiative (see Insel, Landis & Collins, 2013; Koroshetz et al., 2018) places emphasis on identifying specific anatomic and physiologic properties of cells within the brain. In order to do this rigorously and objectively, a systematic way of assigning cells and axonal projection targets to appropriate brain regions is required. Unfortunately, the identification of distinct brain regions in a tissue section under study (thereafter, “experimental section”) with the currently available tools (printed and digital brain atlases) is tedious, cumbersome and prone to error. The difficulties faced by researchers using brain atlases to identify brain regions in experimental sections are illustrated in Figure 1a for the mouse brain through an example from the GENSAT (Gene Expression Nervous System Atlas) project (Gerfen, Paletzki & Heintz, 2013; Pastrana, 2014). Identification of the regions of interest (green and yellow arrows in Fig. 1a) by means of commonly used approaches is hampered by many factors. First, an investigator must move back and forth across plates of a mouse brain atlas to find a section in the atlas that is most similar to the experimental section. Then, the three-dimensional (3D) boundaries of the regions of interest need to be inferred and translated from the static medium of the atlas to the experimental section visualized as an image or under a light microscope. This process must then be repeated for every experimental section under investigation. Second, finding a mouse brain atlas with scale and resolution that complements the experimental section image is complicated by the fact that the cutting angle of histologic sections may deviate from canonical planes. This issue is practically unavoidable when sectioning mouse brains. The process of matching an experimental section image to a plane in a mouse brain atlas when the experimental section is not precisely aligned with the atlas is difficult and results in highly subjective judgments. To date, no readily-available tool exists for researchers to match oblique histology sections to multiple planes of a brain atlas, objectively and automatically.

**FIGURE 1.**
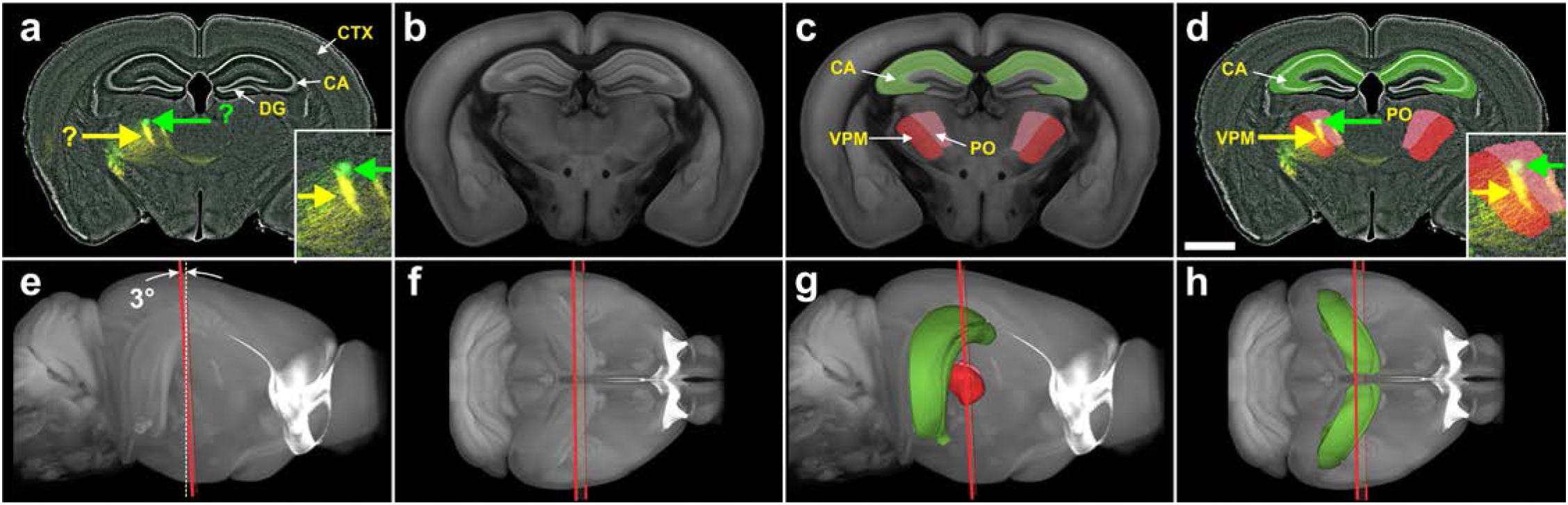
Automatic identification and delineation of regions in a mouse brain section. (**a**) Digital image of a coronal, 80 μm-thick frozen section of the brain from a Cre-driver mouse (from the GENSAT project) in which two fluorescent reporters (green and yellow arrows) label axons from two injection sites (fluorescent Nissl counterstain in gray). In these mice Cre recombinase is expressed under the control of the Sim1(KJ18 line) promoter which labels a subset of layer 5 pyramidal cells. Fluorescent protein expression is driven by stereotaxic injection of adeno-associated virus (AAV) vectors. Stereotaxic injection of AAV vectors into a specific part of the cerebral cortex drives Cre-dependent expression of one or more fluorescent reporters. Projections of these neurons to other subregions of the brain can then be identified by fluorescence microscopy. The counterstain allows investigators with basic knowledge in mouse brain anatomy to identify some key regions (white arrows). However, the identification of the regions of interest (green and yellow arrows) requires expert knowledge in mouse brain anatomy and the consultation of a mouse brain atlas. (**b**) The best fitting oblique angle slice that corresponds to the section shown in a, that was automatically extracted from the 3D reference image of the CCF v3. (**c**) The slice shown in b, with semitransparent overlay of some regions that were automatically extracted from the spatially aligned 3D reference image of the CCF v3. (**d**) The section shown in a, with automatic overlay of the regions shown in c. In this example, the regions of interest were the posterior complex of the thalamus and the ventral posteromedial nucleus of the thalamus. (e-h) Plane of section (red lines) of the slice shown in b through the 3D reference image of the CCF v3, with (g, h) and without (e, f) display of the regions shown in c contained in the spatially-aligned 3D annotations of the CCF v3 (e and g, view from the side; f and h, view from dorsal). The plane of section was tilted at an angle of 3 degrees to the coronal plane of the CCF v3 (white dotted line in e). Abbreviations: CA, CA region of the hippocampus; CTX, cerebral cortex; DG, dentate gyrus of the hippocampus; PO, posterior complex of the thalamus; VPM, ventral posteromedial nucleus of the thalamus. The scale bar in (d) represents 2 mm in (a) and (d) and 1 mm in the high-power insets in (a) and (d).

To overcome the challenges of subjective judgments in atlas referencing, we developed a novel brain navigation system (NeuroInfo) that 1) automatically registers images of experimental mouse brain sections with a 3D digital mouse brain atlas that is essentially based on the third version of the Allen Mouse Brain Common Coordinate Framework (CCF v3) (Allen Institute for Brain Science, 2017); 2) retrieves graphical region delineations and annotations from this 3D digital mouse brain atlas; and 3) superimposes this information onto the experimental section image on a computer screen. Data annotation is essentially achieved in two steps: section-to-atlas registration followed by atlas-based segmentation.

The term 3D digital mouse brain atlas is defined here as a collection of 3D digital files comprising anatomic mouse brain images and associated delineations, which includes the following elements: a 3D reference image embedded in a physical coordinate system, spatially-aligned 3D annotations, and a related ontology that provides the relationship among regions. The 3D reference image of the CCF v3 is based on a 3D, 10 μm isotropic, highly detailed population average of 1675 mouse brains that were imaged with serial two-photon tomography using a TissueCyte 1000 system (TissueVision, Somerville, MA, USA), aligned to a common frame. The 3D annotations of the CCF v3 include delineations of 738 regions based on multimodal references and the related ontology (Allen Institute for Brain Science, 2017).

Here we present a detailed description of the major components of NeuroInfo and their functionality, as well as the results of extensive validation studies performed on 60 experimental coronal sections from 12 different mouse brains that were generated in two different laboratories and imaged with either fluorescence or brightfield microscopy.

## METHODS

### Creation of software and implementation of section-to-atlas registration in NeuroInfo

NeuroInfo was developed as a fully-integrated desktop application written in C++, using an object-oriented design approach.

We used a specialized intensity-based 3D image registration method to establish the alignment of two-dimensional (2D) images of experimental sections (thereafter, “experimental section images”) to the 3D reference image of the CCF v3 (see Brown, 1992). Our implementation is based on the National Library of Medicine Insight Segmentation and Registration toolkit (ITK) with several significant customizations and modifications (Avants et al., 2014). In general, image registration optimizes the parameters of a transform that maps points from a “fixed” image onto corresponding points in a “moving” image.

The optimization process minimizes the value of an image comparison metric that evaluates how well the images align based on intensity values observed at corresponding locations (Zitová and Flusser, 2003). In our implementation, the registration is performed in physical space—an origin and pixel/voxel spacing establishes a coordinate frame for each image—to make the transform independent of the digital resolution of the images involved. The 2D experimental section image serves as the fixed image and is treated as a 3D image that has a thickness on the order of the physical section along the sectioning direction (e.g. 80 μm). Our current implementation supports visualization of the cytoarchitecture in experimental section images with fluorescent (as shown in Fig. 2a-c) or thionin (as shown in Fig. 2d-f) Nissl stains, and these single channels were used for registration. The two-photon autofluorescence reference image of the CCF v3 was used as the moving image.

**FIGURE 2.**
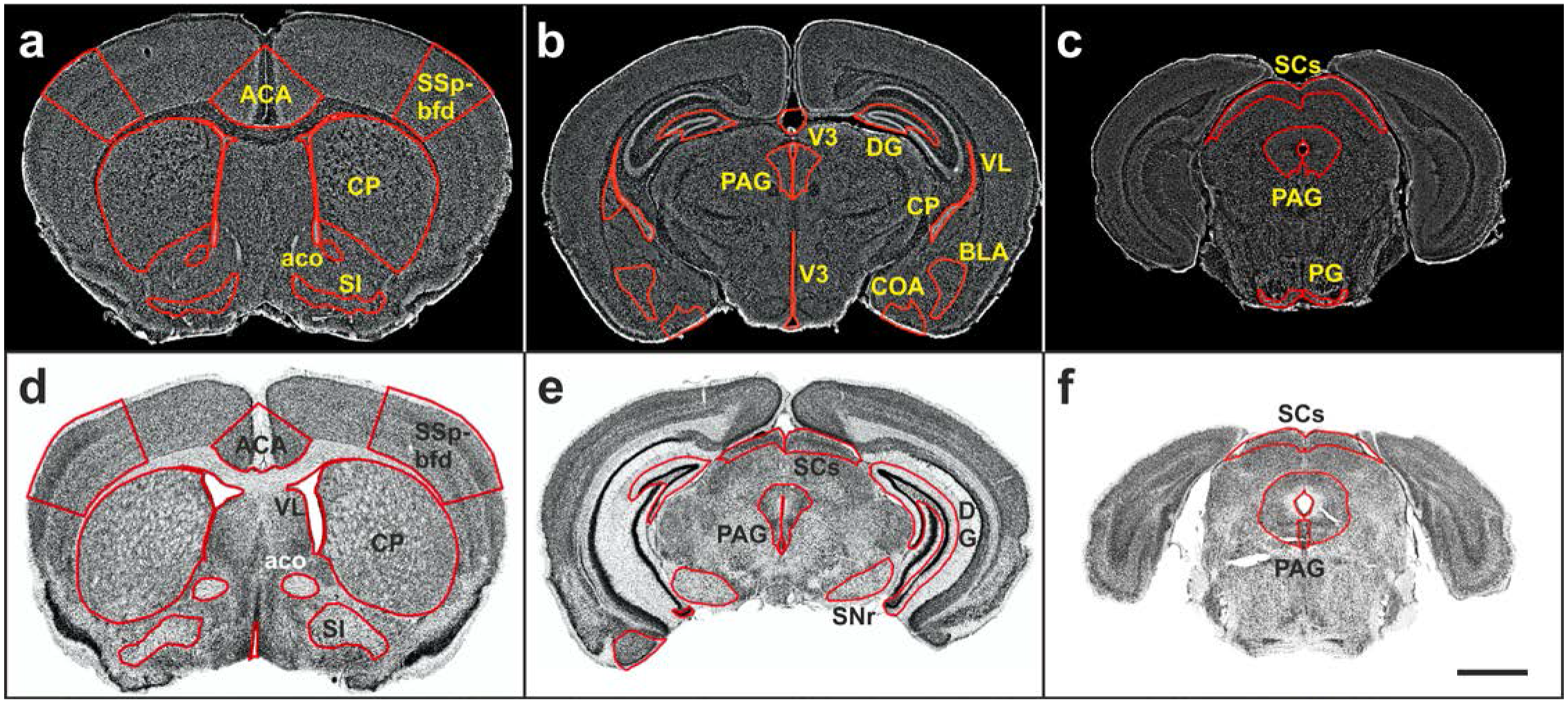
Virtual slides of representative coronal sections (experimental section images) of mouse brains with overlay of delineations of the regions investigated in the present study. (**a-c**) Sections of mouse brains from the GENSAT project. (**d-f**) Sections provided by the CHDI Foundation. Details are provided in the main text. Abbreviations: ACA, anterior cingulate area; aco, anterior commissure (olfactory limb); BLA, basolateral amygdala nucleus; COA, cortical amygdalar area; CP, caudoputamen; DG, dentate gyrus; PAG, periaqueductal gray; PG, pontine gray; SCs, superior colliculus (sensory related); SI, substantia innominata; SNr, substantia nigra (reticular part); SSp-bfd, primary somatosensory area (barrel field); VL, lateral ventricle; V3, third ventricle. The scale bar in (f) represents 2 mm in (a-f).

The specialized image registration process employs multiple stages of multiple-resolution coarse-to-fine refinement. Each stage of the process models an aspect of the physical process of histologic embedding, sectioning and mounting, and operates on a subset of transform parameters. The first stage is initialized with a rigid orientation transform that accounts for the plane of sectioning and the predominant orientation in which the experimental section was imaged (e.g. coronal sections processed along the rostral-caudal axis of the mouse brain with the dorsal surface appearing at the top of the image). The first stage incorporates a search at a low image resolution through the 3D reference image of the CCF v3 along the sectioning axis, using a small number of transform parameters that maintain an orthogonal virtual sectioning through the 3D reference image of the CCF v3. This stage models the position of the experimental section along the sectioning axis and the orientation of the experimental section mounted on a glass slide.

The second stage optimizes the translation, oblique rotation and anisotropic scale of the experimental section image. This stage models the orientation of the tissue block within the microtome and accounts for tissue shrinkage due to histologic processing. The final stage optimizes a nonlinear transform that models deformation of a uniform 3D grid with B-splines. This stage models variability in anatomic structures between individual brains. This multiple stage approach enables exploring the very large transform parameter space in a computationally tractable manner that reflects physical aspects of the problem.

In some cases, the initial search stage does not uniquely identify the location of the experimental section image within the 3D reference image of the CCF v3. Therefore, multiple transform hypotheses can be maintained through all stages of registration. At each stage, transform hypotheses that have a very poor image comparison metric compared to the best hypotheses as well as transform hypotheses that are very similar to higher-ranking transforms, are removed. This approach enables recovery from initially incorrect linear search results.

The image comparison metric serves a critical role in image registration by informing the optimization process how well a particular set of transform parameters aligns the experimental section image with the 3D reference image of the CCF v3. Because the experimental section image was visualized with either fluorescent or thionin Nissl stains and the 3D reference image of the CCF v3 was visualized with two-photon autofluorescence, we selected the Mattes mutual information image comparison metric (Mattes et al., 2001; 2003). We modified the mutual information metric to apply different weights to different parts of the experimental section image preferentially to enforce a better anatomic match. We found that using gradient-based weights of the experimental section image improved the alignment to strong features in the mouse brain, such as white matter tracts.

### Implementation of atlas-based segmentation in NeuroInfo

The transform recovered via image registration enables atlas-based segmentation: the transfer of information from the CCF v3 to the experimental section image or from the experimental section image into the CCF v3, assigning image pixels to brain regions in the reference atlas. The 3D reference image and 3D structural annotations of the CCF v3 and the ontology of the Allen Reference Atlas (thereafter: “ARA ontology”) (Allen Institute for Brain Science, 2017) enable sophisticated queries that provide anatomic context to experimental section images. We developed the following capabilities enabled by section-to-atlas registration:

#### Creation of an atlas image that matches the experimental section image

Image resampling is used to extract 2D atlas images (oblique angle slices) from the 3D reference image of the CCF v3 (Zitová and Flusser, 2003). The coordinates of the desired resampled atlas image are created by duplicating the physical coordinate information from the experimental section image. The forward transform recovered by registration transforms each point in the resampled atlas image into the 3D reference image space of the CCF v3, and linear interpolation of the 3D reference image of the CCF v3 determines the intensity of each pixel in the resampled atlas image. This enables extracting from the 3D reference image of the CCF v3 the best fitting oblique angle slice that corresponds to the experimental section and provides a visualization of the registration accuracy (Fig. 1b).

#### Identification of regions of interest in the experimental section

The investigator identifies a point in the experimental section image using the computer mouse. The forward transform recovered by section-to-atlas registration transforms the point into the coordinate space of the 3D reference image of the CCF v3. The anatomic identity of this point is then retrieved from the 3D annotations of the CCF v3, and the ARA ontology provides the anatomic region name and its relationship to other anatomic regions. This region name is updated on the computer screen in real time (Fig. 1c-d), which enables full exploration of the entire experimental section’s anatomy (Fig. 3). Figure 4 demonstrates that the approach also works when the sectioning angle deviates from canonical planes presented in conventional mouse brain atlases.

**FIGURE 3.**
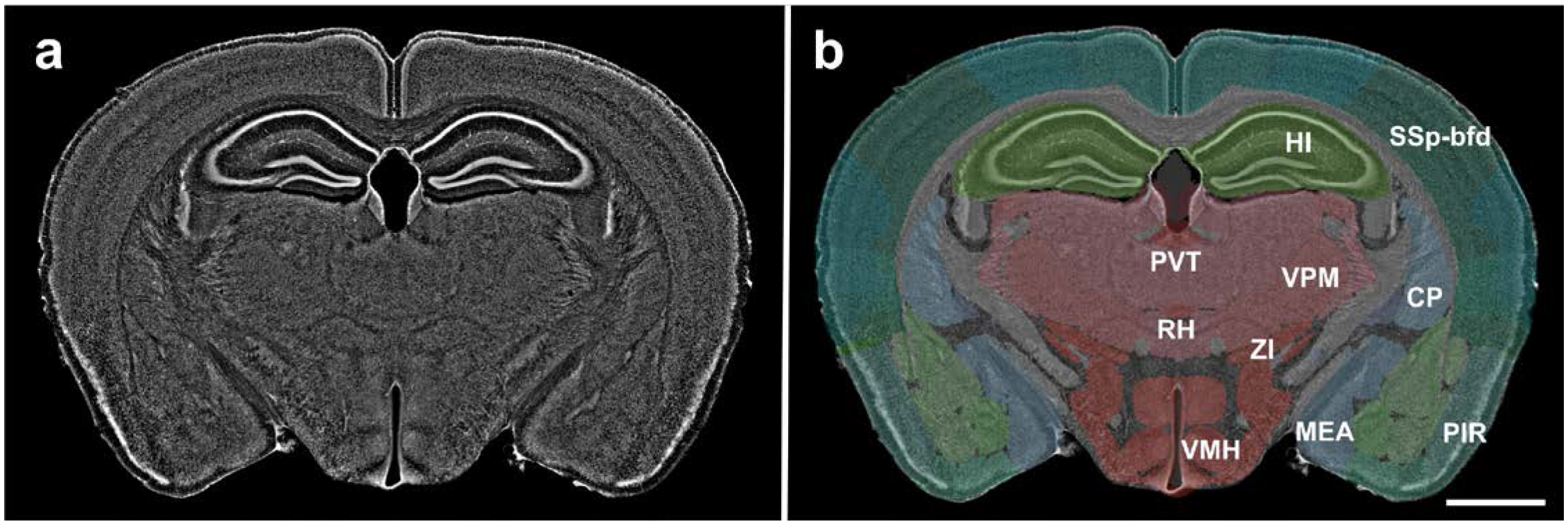
The image shown in Fig. 1a (fluorescent Nissl counterstain shown in gray), with (**b**) and without (**a**) semitransparent overlay of 147 distinct anatomical regions automatically annotated by NeuroInfo (10 of these 147 regions are annotated in this example). Details are provided in the main text. Abbreviations: CP, caudoputamen; HI, hippocampus; MEA, medial amygdalar nucleus; PIR, piriform area; PVT, paraventricular nucleus of the thalamus; RH, rhomboid nucleus; SSp-bfd, primary somatosensory area, barrel field; VMH, ventromedial hypothalamic nucleus; VPM, ventral posteromedial nucleus of the thalamus; ZI, zona incerta. The scale bar in (b) represents 2 mm in (a, b).

**FIGURE 4.**
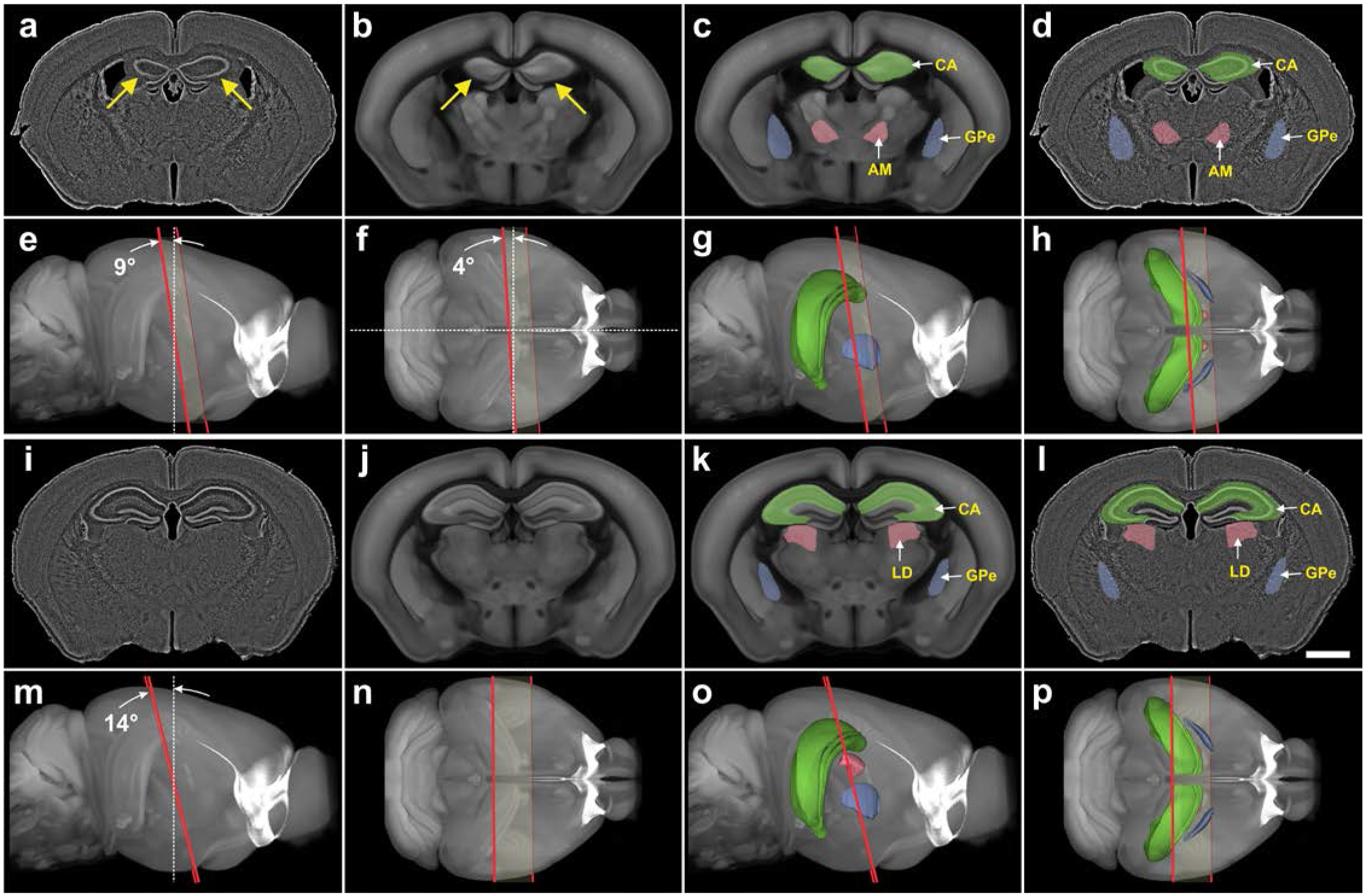
Automatic identification and delineation of regions in mouse brain sections with cutting angles deviating from canonical planes. (**a**, **i**) Digital images of nearly coronal 80 μm thick frozen experimental sections of the brain from a Cre-driver mouse (from the GENSAT project; only fluorescent Nissl counterstain shown in gray). Note the asymmetry of the hippocampus between the left and right brain half (arrows in a). (**b**, **j**) The best fitting oblique angle slices that correspond to the experimental sections shown in a and i, that were automatically extracted from the 3D reference image of the CCF v3. The asymmetry of the hippocampus between the left and right brain half is also found in the slice shown in b (arrows). (**c**, **k**) The slices shown in b and j, with semitransparent overlay of three regions automatically extracted from the spatially aligned 3D reference image of the CCF v3. (**d**, **l**) The sections shown in a and i, with automatic semitransparent overlay of the regions shown in c and k. (**e-h** and **m-p**) Plane of section (red lines) of the slices shown in b and j through the 3D reference image of the CCF v3, with (g, h, o, p) and without (e, f, m, n) display of the regions shown in c and k contained in the spatially aligned 3D annotations of the CCF v3. (e, g, m, o) view from the side; (f, h, n, p) view from dorsal. The plane of section of the slice shown in b was tilted at an angle of 9 degrees from the dorsal-ventral axis (white dotted line in e) and 4 degrees from the lateral axis (white dotted line in f) with respect to the coronal plane of the CCF v3. The plane of section of the slice shown in j was tilted at an angle of 14° to the coronal plane of the CCF v3 (white dotted line in m). Nevertheless, in both cases it was possible to automatically identify and delineate correctly regions in the experimental sections. Abbreviations: AM, anteromedial nucleus of the thalamus; CA, CA region of the hippocampus; GPe, globus pallidus, external segment; LD, lateral dorsal nucleus of the thalamus. The scale bar in l represents 2 mm in a, d, i, l.

Determination of the boundaries of a region in the experimental section image – The investigator uses the computer mouse to select a region of interest in the experimental section image or queries a region from the ARA ontology. An annotation image that matches the experimental section image is created using the same resampling process described above, but using nearest-neighbor interpolation of the 3D annotations of the CCF v3 in the final step. The ARA ontology provides the unique identification number of the region of interest and all contained sub-regions. This information is used to build a mask for the region of interest within the resampled annotation image. The connected components algorithm identifies and traces the boundaries of each distinct region in the mask (Shapiro and Stockman, 2001) (Fig. 1d). The result is that the anatomic delineations become part of the data that can be used during analysis of the experimental section.

### Visualization of the CCF v3 in 3D

For visualization purposes, we built smooth anatomic 3D models for each region listed in the ARA ontology based on the 3D annotation image of the CCF v3. This was achieved using the following procedure. For each anatomic region, a 3D image mask was created from the 3D annotation image of the CCF v3 using the union of the anatomic region and all contained sub-regions determined from the ontology.

A 3D triangular mesh was created using the marching cubes algorithm (Lorensen and Cline, 1987). The mesh was smoothed and decimated to reduce the number of triangles in the model (c.f. Garland and Heckbert, 1997).

The mesh models were saved to disk and later recalled for 3D visualization. The 3D reference image of the CCF v3 is visualized using GPU-accelerated volume rendering with maximum intensity projection (e.g. Fig. 1e). Selected anatomic regions are visualized by loading the constructed 3D meshes and displaying them within the volume rendered 3D reference image of the CCF v3 (e.g. Fig. 1g). The location of the sectioning plane of the experimental section image is visualized by transforming a planar region that encloses the experimental image into the 3D reference image’s space using the linear components of the section-to-atlas registration transform (e.g. Fig. 1e-h).

### Validation of NeuroInfo

A total of 461 coronal, 80 μm-thick frozen serial sections from 78 adult male FVB/N/CD1 mouse brains (from the GENSAT project) were stained with a fluorescence Nissl stain (Neurotrace Blue; Invitrogen, Carlsbad, CA, USA) and imaged using a AxioImager Z1 microscope (Carl Zeiss Microscopy, Jena, Germany) with a 10× objective (N.A. = 0.30, a motorized stage (Ludl Electronic Products, Hawthorne, NY, USA), and a Hamamatsu Orca Flash 4 camera (Hamamatsu City; Japan) controlled by Neurolucida software (MBF Bioscience, Williston, VT, USA) to generate 2D virtual slides (thereafter: “GENSAT section images”). Each GENSAT section image contained between 80 to 200 tiles. Representative examples are shown in Fig. 2a-c.

Another set of 2D virtual slides was generated from 691 coronal 60 μm-thick serial sections of 21 brains from adult male and female C57/BL6 mice that were provided by the CHDI Foundation (New York, NY) in the framework of a research project commissioned to MBF Bioscience (thereafter: “CHDI section images”). These sections were stained with thionin and imaged with a slide scanner equipped with a 20× objective (N.A. = 0.75; BLiSS-200, MBF Bioscience, Williston, VT, USA). Again, each CHDI section image contained between 80 to 200 tiles. Representative examples are shown in Fig. 2d-f.

During pilot experiments, we determined that the registration approach depended on finding a good initial position of the experimental section image within the 3D reference image of the CCF v3. Specifically, a linear search along the sectioning axis worked well when dealing with whole experimental section images of good quality (such as those shown in Fig. 2), but it required manual initialization when dealing with images of partial or damaged experimental sections. Sections were manually inspected for histology artifacts such as large tears, folds and missing pieces. Sections with significant artifacts were omitted. A random subset of 30 GENSAT section images and 30 CHDI section images were then selected for testing. These 60 experimental section images from 12 different mouse brains were automatically registered by means of nonlinear registration with the 3D reference image of the CCF v3 using NeuroInfo. We recorded the X- and Y-scale components of the transform, Mattes mutual information metric (Mattes et al., 2001; 2003), and processing time during registration.

Then, nine GENSAT section images and six CHDI section images were randomly selected, and 14 regions were automatically identified and delineated using NeuroInfo. These 14 regions represent different cortical and subcortical regions in the rostral, caudal, middle, lateral, dorsal and ventral parts of the mouse brain (c.f. Fig. 2a-f). If a region did not exist in an experimental section image, NeuroInfo did not draw any delineations upon region selection.

The same regions were manually delineated by a technician on the 15 selected experimental section images using the Stereo Investigator software (MBF Bioscience), referencing the 3D reference image of the CCF v3 and the ARA ontology. Then, the 15 experimental section images overlain with both the manual delineations (thereafter: “D_m_s”) and the automatic delineations (thereafter: “D_a_s”) were checked by neuroanatomy experts (CRG, PRH), who annotated problematic or inappropriate Dms and Das. Inappropriate Dms were corrected.

After correction of inappropriate D_m_s, differences between D_a_s and the corresponding D_m_s were quantified by means of the following metrics: 1) overlap via the Dice coefficient, calculated as

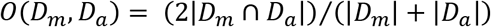

(Dice, 1945; Fischl et al., 2002; Klein et al., 2009); 2) the distance between the centroids of D_a_ and D_m_; 3) the relative number of D_a_s in which the centroid of D_a_ was found both within D_a_ and D_m_; 4) the difference between the areas of D_a_ and D_m_, calculated as

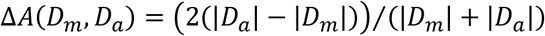

(Fischl et al., 2002; Klein et al., 2009); 5) the percentage of area-D_m_ that was covered by area-D_a_, calculated as

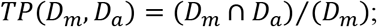

and 6) the amount of area-D_a_ outside area-D_m_ expressed as a percentage of area-D_m_, calculated as

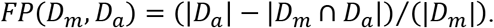

### Statistical analysis

Mean and standard deviation of the X- and Y-scale components of the transform, the values obtained by calculating the Mattes mutual information metric and the processing time were separately calculated for the 30 GENSAT section images and the 30 CHDI section images. The Shapiro-Wilk normality test was used to determine whether the distributions of these data were consistent with a Gaussian distribution. Differences between groups were tested with two-way ANOVA (X- and Y-scale components of the transform) and the Mann Whitney test (values obtained by calculating the Mattes mutual information metric and processing time). In all analyses, an effect was considered statistically significant if its associated p value was smaller than 0.05. All calculations were performed using GraphPad Prism (version 5.00 for Windows, GraphPad Software, San Diego, CA, USA).

## RESULTS

### Major components of NeuroInfo and their functionality

The Graphical User Interface (GUI) of NeuroInfo and its Visualization and Navigation Tool is shown in Figure 5 (the details in parentheses refer to the specific example shown in this figure). In addition to the navigation bar (“a” in Fig. 5) and toolbar (“b” in Fig. 5) for basic operations (e.g., opening images, saving and loading registration results, etc.) the major components of section registration are a Macro View window (“c” in Fig. 5) which shows an experimental section image under investigation (here a GENSAT section image), an Ontology window (“d” in Fig. 5) that displays anatomical structures in a hierarchical tree and allows users to select regions for automatic delineation (here the primary somatosensory area, barrel field [SSp-bfd]), a zoomable, full-resolution Main Image window (“e” in Fig. 5) displaying a high-power view of the selected region(s) (here SSp-bfd) in the experimental section image overlain by the region’s delineation automatically generated by NeuroInfo’s Registration Match Finding and Identification Tool (conceptually described in Figs 1 and 3), a Reference Slice window (“f” in Fig. 5) showing the oblique angle slice through the 3D reference image of the CCF v3 that best fits the experimental section image, with semitransparent overlay of the selected region(s) (here the SSp-bfd), a 3D Reference Image window (“g” in Fig. 5) showing the best fitting oblique angle slice (red line) through the 3D reference image of the CCF v3 (seen from the side), displaying the region(s) selected in the Ontology window (here SSp-bfd), and a Section Registration window (“h” in Fig. 5) that allows the user to control section registration using automated linear or nonlinear registration, or by fine-tuning section registration by adjusting translation, rotation and scaling. The NeuroInfo Atlas Calibration Tool (Fig. 6) enables calibration of the X and Y scaling to correct for tissue compression caused by histologic processing, sectioning and mounting. For doing so the user selects 1) the imaging modality (fluorescence or brightfield), and in the case of fluorescence imaging, the fluorescence registration channel (Fig. 6a), 2) the anatomic sectioning plane (Fig. 6b), 3) the sectioning direction (in the example shown in Fig. 6c ‘Rostral-most cut first’ was selected, indicated by the blue frame around the panel on the right), and 4) what is at the bottom of the experimental section image (in the example shown in Fig. 6d ‘Ventral’ was selected, indicated by the blue frame around the panel on the top left). The Atlas Calibration Tool then guides the user through the interactive registration of a single experimental section image to determine the relative scaling between the experimental section image and the 3D reference image of the CCF v3. Our validation experiments determined that this laboratory-specific scaling calibration was critical to enable successful automatic registration of images of novel experimental sections that were processed in the same manner (details are provided in Fig. 7). Once completed, the scaling calibration is used for subsequent section registration operations.

### Registration of experimental section images with the 3D reference image of the CCF v3

In order to match the 60 selected experimental section images as closely as possible with the 3D reference image of the CCF v3 it was necessary to adjust the scaling in X on average by 1.09 (GENSAT section images) and 1.12 (CHDI section images), and in Y on average by 1.21 (GENSAT section images) and 1.27 (CHDI section images). Both the between-groups differences (GENSAT section images vs. CHDI section images) and the within-groups differences (X-scaling vs. Y-scaling) were statistically significant (two-way ANOVA; pLab < 0.001; pDirection < 0.001) (Fig. 7a).

The mean value obtained by calculating the Mattes mutual information metric was 0.37 (GENSAT section images) and 0.16 (CHDI section images). This difference was statistically significant (Mann Whitney test; p < 0.001) (Fig. 7b).

The average processing time for registering the images of the experimental sections was 82 s (GENSAT section images) and 72 s (CHDI section images), with no statistically significant difference between the groups (Mann Whitney test; p = 0.819) (Fig. 7c).

**FIGURE 5.**
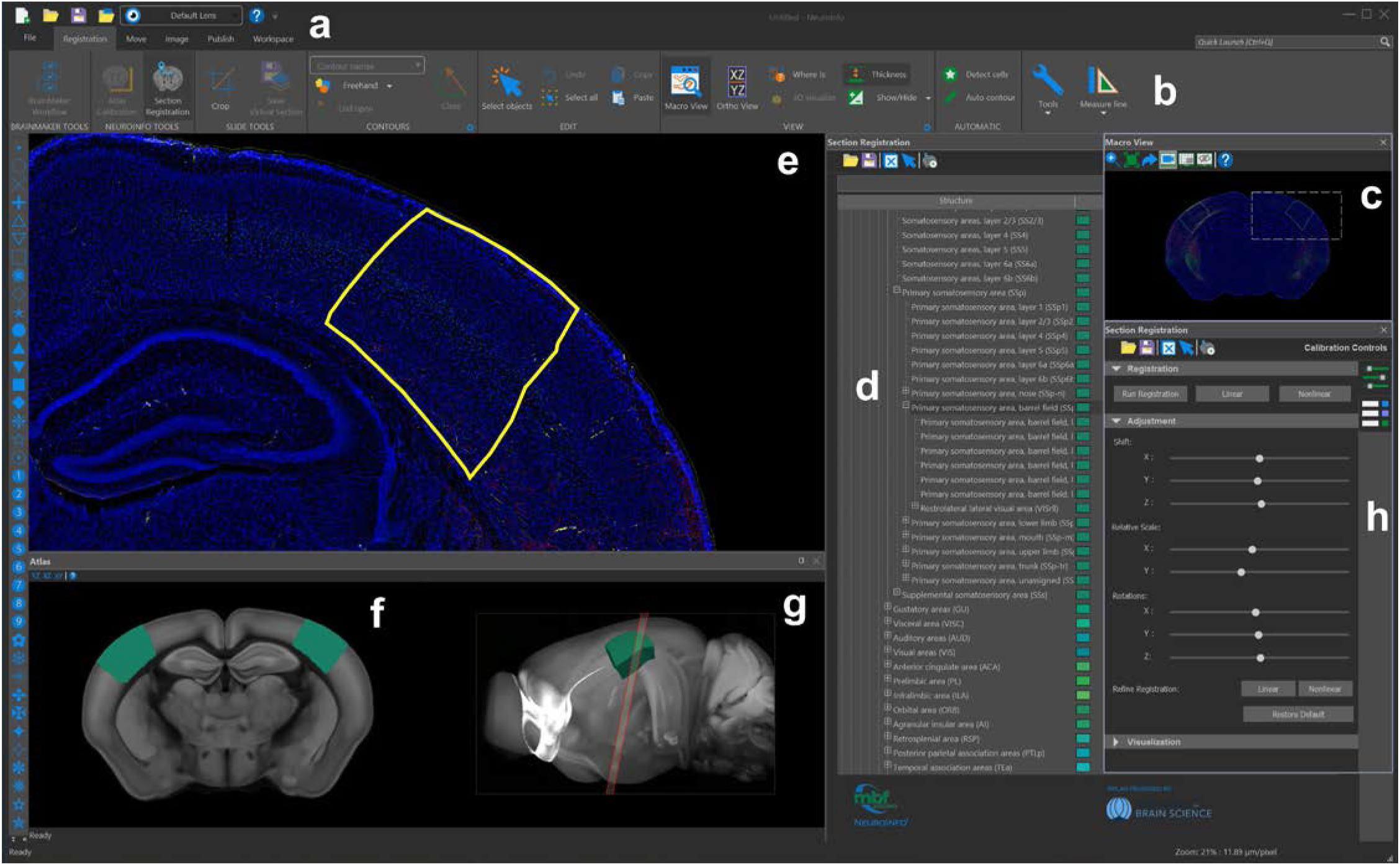
The graphical user interface of NeuroInfo. The navigation menu (**e**) displays a sample image in the coronal plane, with barrel cortex structures selected in yellow and annotations in teal (**d**, ontology; **f**, **g**, reference space). (**a**, **b**, **h**): control toolboxes and viewers.

### Identification of regions of interest in experimental section images

For the regions caudoputamen (CP), dentate gyrus (DG), primary somatosensory area (barrel field) (SSp-bfd), substantia nigra (reticular part) (SNr), anterior cingulate area (ACA), periaqueductal gray (PAG) and superior colliculus (sensory related) (SCs) (green frames in Fig. 8a; green bars in Fig. 8b-h) excellent alignments between D_a_ and D_m_ were obtained, indicated by mean Dice coefficients >0.7 (Fig. 8b), average distances between the centroids of D_a_ and D_m_ <200 μm (Fig. 8c), relative numbers between 83% and 100% of D_a_s in which the centroid of D_m_ was found within D_a_ (Fig. 8d), absolute mean differences between the areas of D_a_ and D_m_ <0.1 (except SCs) (Fig. 8e), mean TP values >75% (except SCs) (Fig. 8f) and mean FP values <50% (Fig. 8g). Figure 8h demonstrates that among these regions were the largest ones that were investigated. Furthermore, Figure 8a shows (with the exception of SNr) that these regions are located in the dorsal parts of the mouse brain.

For the regions basolateral amygdala nucleus (BLA), substantia innominata (SI) and cortical amygdalar area (COA) (yellow frames in Fig. 8a; yellow bars in Fig. 8b-h) the alignments between D_a_ and D_m_ were less precise, indicated by mean Dice coefficients between 0.5 and 0.7 (Fig. 8b). This was supported by all other metrics that returned results that were not as good as in case of the regions CP, DG, SSp-bfd, SNr, ACA, PAG and SCs (Fig. 8c-g). Figure 8h demonstrates that the regions BLA, SI and COA were among the smallest ones that were investigated. Furthermore, Figure 8a shows that these regions are located in the ventrolateral parts of the mouse brain.

For the regions pontine gray (PG), anterior commissure (olfactory limb; aco), lateral ventricle (VL) and third ventricle (V3) (red frames in Fig. 8a; red bars in Fig. 8b-h) the alignments between D_a_ and D_m_ were poor, indicated by mean Dice coefficients <0.5 (Fig. 8b).

**FIGURE 6.**
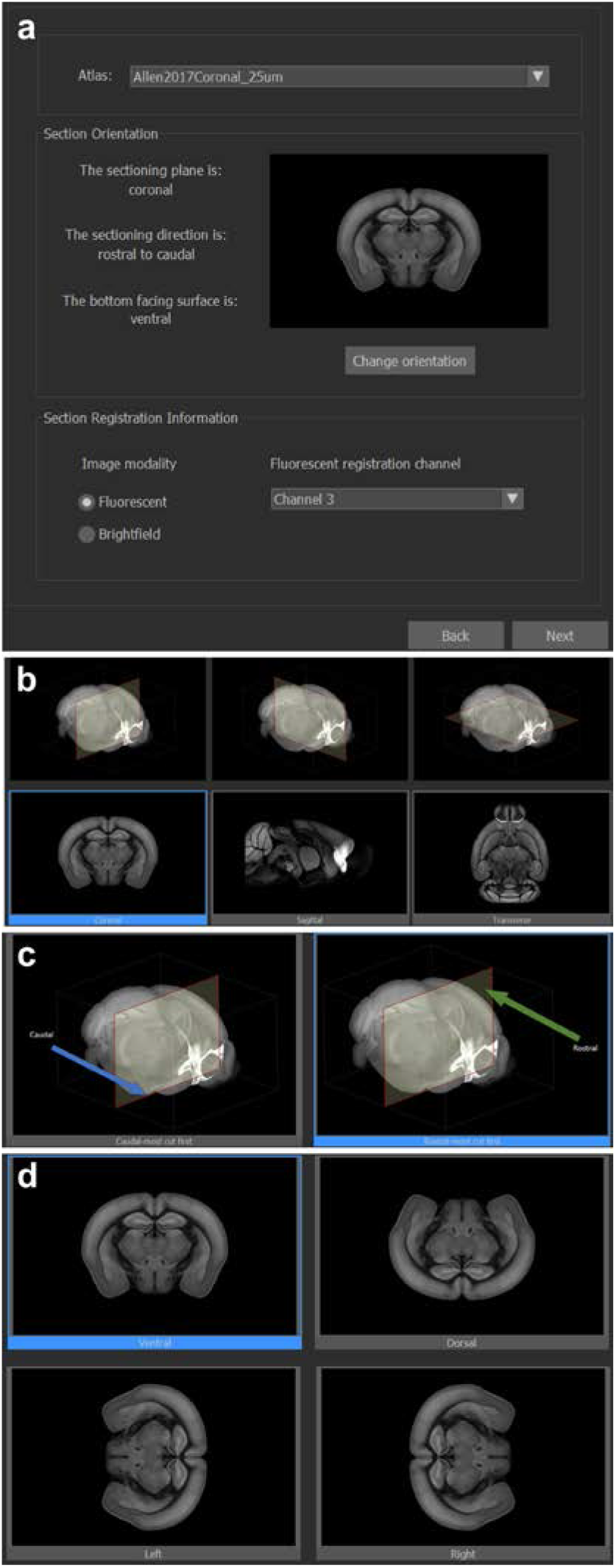
The NeuroInfo Atlas Calibration Tool and anatomic orientation selection interface. (**a**) atlas and 2D section image specification; (**b**) anatomic sectioned plane selection; (**c**) sectioned direction selection (coronal); (**d**) section image orientation selection (coronal).

The results obtained with the other metrics (except FP) were even poorer than those obtained for the regions BLA, SI and COA (Fig. 8c-g). Figure 8h demonstrates that the regions PG and aco were the smallest ones that were investigated. Figure 8a shows that these regions are located in the middle parts of the mouse brain.

**FIGURE 7.**
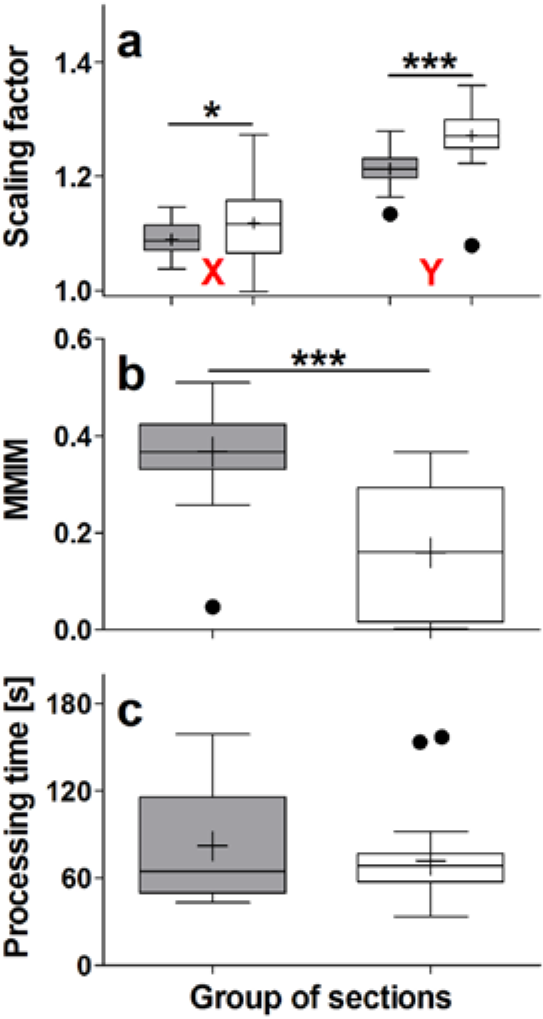
Results of automatically registering 30 GENSAT section images (gray bars) and 30 CHDI section images (open bars) investigated in the present study with the 3D reference image of the CCF v3, using NeuroInfo. The panels show Tukey boxplots of (**a**) the scaling factors in X and Y, (**b**) the values obtained by calculating the Mattes mutual information metric (MMIM), and (**c**) the processing time, in seconds. Statistically significant differences between the groups are indicated in panels a and b (*; p < 0.01; ***, p < 0.001).

## DISCUSSION

The results of the validation experiments performed on 60 experimental coronal sections from 12 different mouse brains that were generated in two different labs and imaged with respectively fluorescence or brightfield microscopy can be summarized as follows: 1) NeuroInfo performed remarkably well in automatically delineating regions that are large and/or located in the dorsal parts of mouse brains (mean Dice coefficients >0.7), returned less precise delineations of regions that are small and/or are located in the ventrolateral parts of mouse brains (mean Dice coefficients between 0.5 and 0.7), and must be further improved for the delineation of regions that are respectively either small and located in the middle parts of mouse brains or do not contain nerve cells (mean Dice coefficients <0.5).

**FIGURE 8.**
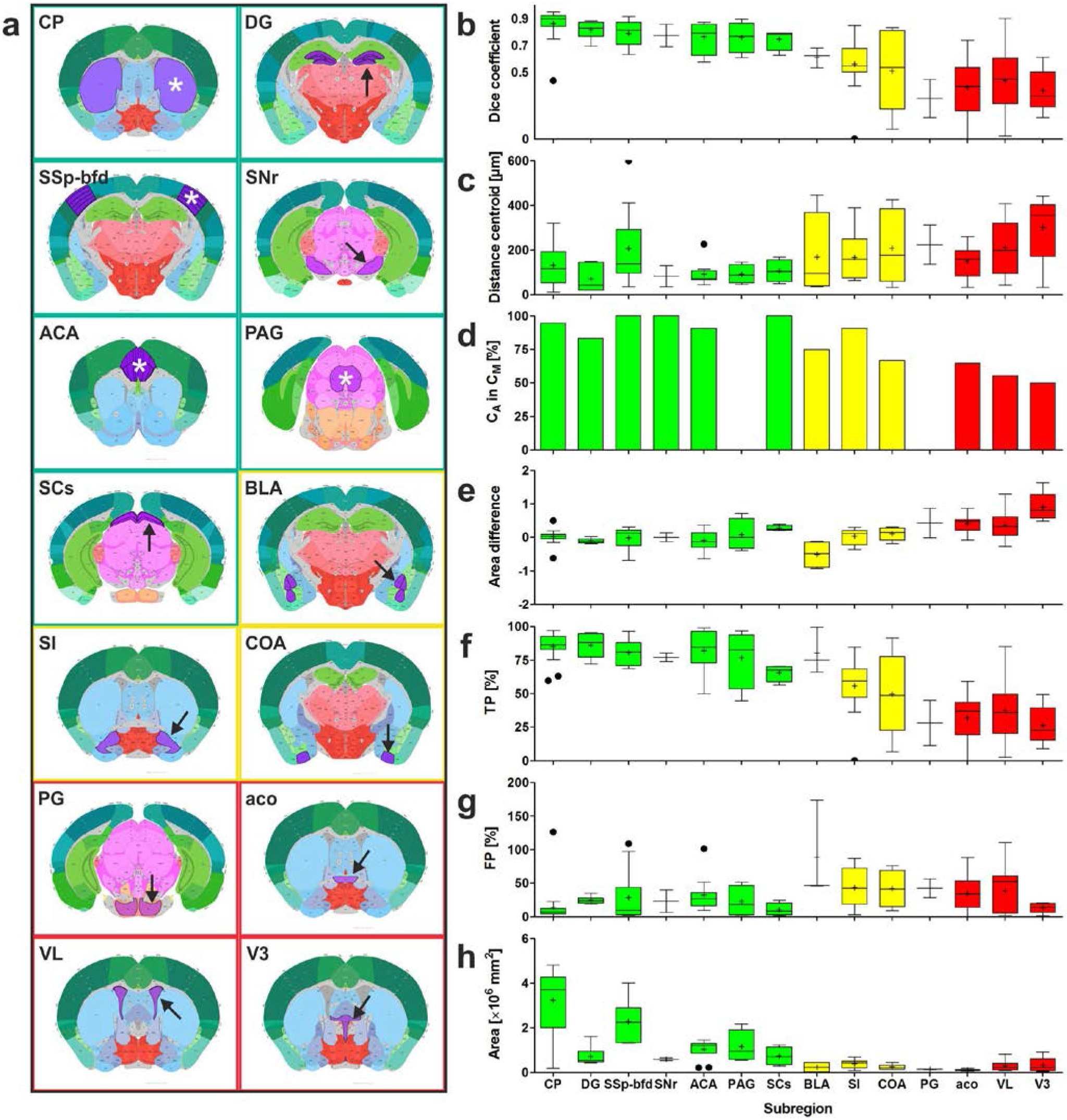
(**a**) Plates of 2D view of the CCF v3 showing those regions (marked by asterisks and arrows) that were investigated during the validation experiments. The colors of the frames match the colors of the bars in (b-h). (**b**-**h**) Results of the validation experiments, comparing manual delineations (D_m_s) performed by neuroanatomy experts (ground truth) with automatic delineations (D_a_s) performed by NeuroInfo of the regions shown in (a) on 15 experimental section images of mouse brains. (b,c,e-h) Tukey boxplots of (b) the Dice coefficient, (c) the distance between the centroids of D_a_ and D_m_, (e) the difference between the areas of D_a_ and D_m_, (f) the percentage of area-D_m_ that was covered by area-D_a_ (TP), (g) the amount of area-D_a_ outside area-D_m_ expressed as a percentage of area-D_m_ (FP), and (h) the absolute area of D_m_. (d) Relative number of D_a_s in which the centroid of D_a_ was found within both D_a_ and D_m_. Green, yellow and red bars indicate those regions for which the Dice coefficient was found to be >0.7, 0.5-0.7, <0.5, respectively. Abbreviations: CP, caudoputamen; DG, dentate gyrus; SSp-bfd, primary somatosensory area (barrel field); SNr, substantia nigra (reticular part); ACA, anterior cingulate area; PAG, periaqueductal gray; SCs, superior colliculus (sensory related); BLA, basolateral amygdala nucleus; SI, substantia innominata; COA, cortical amygdalar area; PG, pontine gray; aco, anterior commissure (olfactory limb); VL, lateral ventricle; V3, third ventricle.

**FIGURE 9.**
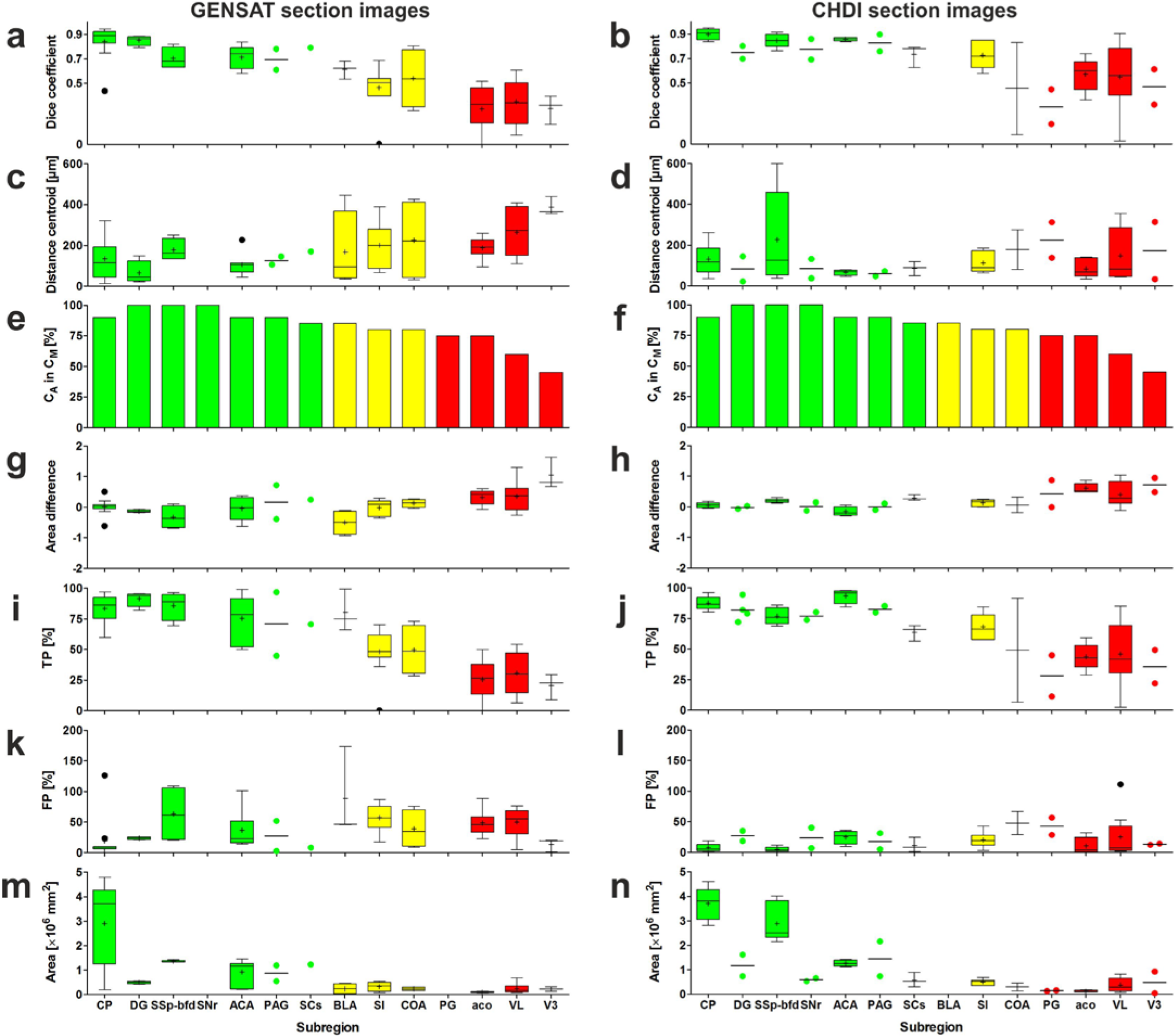
Results of the validation experiments, comparing manual delineations (D_m_s) performed by neuroanatomy experts (ground truth) with automatic delineations (D_a_s) performed by NeuroInfo of the regions shown in Figure 8a on nine experimental section images of mouse brains from the GENSAT project (panels on the left) and on six experimental section images of mouse brains provided by the CHDI foundation (panels on the right). The panels show Tukey boxplots of (**a**,**b**) the Dice coefficient, (**c**,**d**) the distance between the centroids of D_a_ and D_m_, (**g**,**h**) the difference between the areas of Da and Dm, (**i j**) the percentage of area-D_m_ that was covered by area-Da (TP), (**k**,**l**) the amount of area-D_a_ outside area-D_m_ expressed as a percentage of area-D_m_ (FP), and (**m**,**n**) the absolute area of D_m_. If for a certain region only two values were available, the Tukey boxplot was replaced by a scattered dot plot of the same color with a line at the mean. (**e**,**f**) Relative number of D_a_s in which the centroid of D_a_ was found within both D_a_ and D_m_. Green, yellow, and red bars indicate those regions for which the Dice coefficient was found to be >0.7, 0.5-0.7, <0.5, respectively. Abbreviations: CP, caudoputamen; DG, dentate gyrus; SSp-bfd, primary somatosensory area (barrel field); SNr, substantia nigra (reticular part); ACA, anterior cingulate area; PAG, periaqueductal gray; SCs, superior colliculus (sensory related); BLA, basolateral amygdala nucleus; SI, substantia innominata; COA, cortical amygdalar area; PG, pontine gray; aco, anterior commissure (olfactory limb); Vl, lateral ventricle; V3, third ventricle.

The latter is currently ongoing in our labs by implementing in NeuroInfo a novel image registration approach that uses model information from the 3D annotation image of the CCF v3 to influence global and local components of the section-to-atlas transform. 2) To our surprise, NeuroInfo was not able to correctly identify the olfactory limb of the anterior commissure with the metrics used in this study, despite the fact that it can be found easily by eye. One potential reason is that alignment information in other parts of the experimental section image dominated (e.g. strong cortical or subcortical features), compromising the local alignment of the anterior commissure. The model-based global-local registration approach currently under research is expected to improve this situation. 3) The performance of NeuroInfo in automatically delineating regions in sections of mouse brains did not depend on whether the experimental sections were imaged with fluorescence or brightfield microscopy. 4) To match the experimental section images of mouse brains provided by two different laboratories with the 3D reference image of the CCF v3 optimally, it was necessary to adjust the scaling in Y to a significantly greater extent than the scaling in X. Thus, the investigated mouse brains appeared ‘flattened’ compared to the 3D reference image of the CCF v3. The reason for this phenomenon is unknown, but may be due to cutting blade pressure on the brains along the Y axis or differences in histologic processing techniques. 5) The average processing time of registering an experimental section image of a mouse brain with the 3D reference image of the CCF v3 was 77 seconds. After registration, automatic delineation of any region contained in the experimental section image is obtained in real-time when selecting a certain region in the Ontology window of NeuroInfo’s GUI (“d” in Fig. 5). In contrast, neuroanatomists would need between several seconds (for a single region such as the caudoputamen) and a full day (e.g., for all 147 distinct regions contained in the experimental section shown in Fig. 1a) to complete the same task (see Fig. 3).

This novel automatic navigation system may fundamentally change the way researchers investigate sections of mouse brains under a light microscope, similar to the way surgical navigation systems have influenced neurosurgery over the past three decades (Mezger, Jendrewski and Bartels, 2013). Specifically, the approach presented here enhances the validity, reliability and reproducibility of studies that depend on the exact identification of distinct regions in the mouse brain, enables navigation through the complex anatomy of the mouse brain without requiring prior in-depth training in neuroanatomy, offers investigators the potential to analyze many more brain regions than is currently possible, and provides investigators with the means to directly correlate their experimental specimens with a number of reference tools such as GENSAT or the Allen Mouse Brain Connectivity Atlas (Oh et al., 2014). Likewise, the same principle enables an investigator to collate data from multiple subjects across multiple experiments by transforming measurements into a common reference space. This is a particularly valuable feature, given the emphasis on use of shared reference frameworks for collaborative initiatives such as the NIH BRAIN initiative (Insel et al., 2013).

The novel automatic navigation system presented here is currently only tested for sections from mouse brains counterstained with fluorescent or thionin Nissl stains. Future work will extend this technology to accommodate other counterstains (such as DAPI or immunohistochemical markers). Ultimately, this technology is not limited to mouse, and can be applied to histologic sections from other species with sufficient reference data available, such as rat (using the Waxholm space; Papp et al., 2014), non-human primates (Reveley et al., 2017), and ultimately humans for which the Allen Human Brain Atlas (Hawrylycz et al., 2012; Ding et al., 2016) already provides an on-line high-resolution annotated reference map.

## ACKNOWLEDGEMENT

We would like to thank Dr. Amy Bernard (Allen Institute for Brain Science, Seattle, WA, USA) for facilitating the licensing of the Allen Mouse Brain CCF v3 data, and Dr. David Howland (CHDI Foundation/CHDI Management, Princeton, NJ, USA) for providing the opportunity to use the CHDI sections in this study. We also thank Alissa Wilson and Julie McMullen, Isabelle Angstman, Maddie Turneau and Aidan Sullivan (MBF Bioscience) for skillful technical assistance.

## AUTHOR CONTRIBUTIONS

SJT, BSE, NOC, CS and JRG initiated the project. BSE and NOC wrote the software. QW, LN, and DF provided expertise on navigating, interpreting, and 3D modeling with the CCF v3. CRG and BMH provided experimental image data and consulted on software feature development. SJT, CS and JRG designed the validation experiments. SJT, BSE, NOC, CRG, PRH and JRG performed the validation experiments. SJT, BSE, NOC, CRG, PRH, CS and JRG analyzed the data. SJT, BSE, NOC and CS wrote the manuscript. QW, LN, BMH, CRG, PRH and JRG contributed to reviewing and editing of the manuscript.

